# Cell size statistics in cell lineages and population snapshots with different growth regimes and division strategies

**DOI:** 10.1101/2020.05.15.094698

**Authors:** Niccolò Totis, César Nieto, Armin Küper, César Vargas-García, Abhyudai Singh, Steffen Waldherr

## Abstract

Growing populations of bacteria control their growth and division reaching narrow distributions of cell-sizes. In this paper we explored how different combinations of growth regimes and division mechanisms lead to different cell-size statistics in these populations. Deterministic and stochastic modeling were used to describe the size distribution of a population of cells that is observed from two different perspectives: as single cell lineages, i.e. random paths in the lineage tree, or as snapshots, at given times, of a population in which all descendants of a single ancestor cell are observed. Our time-dependent approaches allowed us to obtain both the transient dynamics and the steady state values for the main statistical moments of the cell-size distribution. Also, we established mathematical relationships among the statistics in the two considered perspectives, thus improving our knowledge of how cells control their growth and proliferation.

## I. INTRODUCTION

Bacteria are able to regulate growth and division in order to maintain their size within a defined range, a state defined as *homeostasis*[1], [2]. Traditionally, experimental studies on cell division observe a cell population from two main alternative perspectives[3]. Either they (i) track *single lineages* as observed in the mother machine[4], either (ii) they take *population snapshots* of descendant cells of a common ancestor, like in flow cytometry[5] experiments. Understanding the differences between these experimental viewpoints [6] has become an expanding area of research[7], [8], [9], [10].

In the present work deterministic and stochastic computational modeling approaches were used to represent these two perspectives and to analyze how the heterogeneity of a cell population with respect to cell size evolves as a result of growth and symmetric division. To observe single lineages, at each cell division the model keeps track of only one newborn cell. Consequently, the total cell number of the considered sub-population remains constant over the simulated time. To observe population snapshots, instead, the model tracks all progenies and thus the evolution of the whole population over time. In our analysis we considered some scenarios of experimental relevance characterized by different growth regimes and division mechanisms, we estimated the dynamics of the main statistical properties of the size distribution and compared the obtained results.

In Section II, we introduce the two modeling frameworks we used to describe size dynamics: the Population Balance Equation (PBE), a population-based deterministic approach, and Monte Carlo simulations, using a modified stochastic simulation algorithm. Furthermore, we present how these were exploited to represent the considered scenarios of growth and division. Details of the analytical derivations as well as of the numerical solutions are provided. In Section III we present how four biologically relevant scenarios characterize the components of the models. The comparison of the results obtained with the different modeling approaches is portrayed with figures and further discussed.

## II. METHODS

### A. Modeling frameworks

#### a) Population-based deterministic modeling: the population balance equation (PBE)

In general, the framework of PBEs describes the evolution of particulate systems with respect to an extensive property, like size, mass or molecule counts [11]. Here, PBEs allow us to represent the distribution of cell sizes *s* in the population as a number density function *n*(*t, s*) (or NDF) and to study how it changes over time as a result of the processes of cell growth and cell division.

The dynamics of *n*(*t, s*) is given by the following integro-partial differential equation nnnnnnnn:

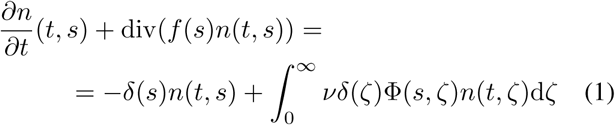

The second term on the left-hand side, the divergence term, describes how the NDF moves through the cell state space. Here, 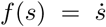 describes the single-cell growth rate, *ν* denotes the number of offsprings generated from each cell division, *δ*(*s*) is the division rate function (in PBE literature, this is often referred to as the breakage rate) and Φ(*s, ζ*) is the kernel function, which defines the probability that the division of a mother cell of size *ζ* generates a daughter cell of size *s*. Φ(*s, ζ*) satisfies the property ∫ Φ(*s, ζ*)d*s* = 1. In our analysis, these functions are defined as follows:

- *ν*. We set *ν* = 1 when, at each division, one single descendant is tracked. We set *ν* = 2 otherwise.
- *f* (*s*). The single-cell growth rate is modeled as *f* (*s*) = *gs*^*a*^, *a* ≥ 0. A linear growth regime is obtained by setting *a* = 0, while, for *a* = 1, cells grow exponentially. A general *a >* 0 can be used to explore other division mechanisms[12].
- *δ*(*s*). The division rate is modeled as *δ*(*s*) = *ks*^*b*^, *b* ≥ 0. In this paper, we limit our analysis to *b* ∈ ℕ. The division rate is thus constant for *b* = 0, size-dependent otherwise.
- Φ(*s, ζ*). The Dirac-delta function is used here to model an equal repartitioning of volume after cell division, so that Φ(*s, ζ*) = 2*δ*(2*s* − *ζ*).

In this way (1) becomes:

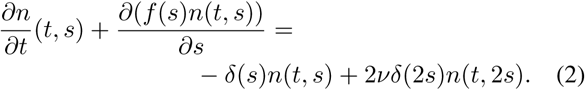

In our analysis, the evolution of the cell size distribution is described through the changes in its raw moments. Starting from the PBE in (1), we provide here an analytical derivation of the moment dynamics. The *i*^*th*^ raw and central moments are denoted as 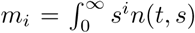 and 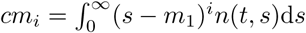 respectively. In the following derivations, we assume that all raw moments remain finite at finite times, which implies lim_*s*→+∞_ *s*^*i*^*n*(*t, s*) = 0. While moments *m*_*i*_(*t*) are always considered as time-dependent variables, we will make use of *m*_*i*_ instead, so that notations are simplified.

The total number of cells at a given time is equivalent to the zeroth moment *m*_0_ of the NDF:

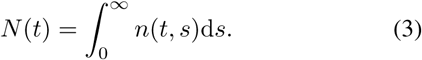

Introducing the normalized NDF

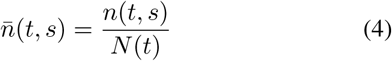

the dynamics of *N* (*t*) can be written as

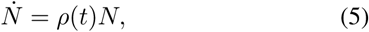

with *ρ*(*t*), the average specific proliferation rate, defined as

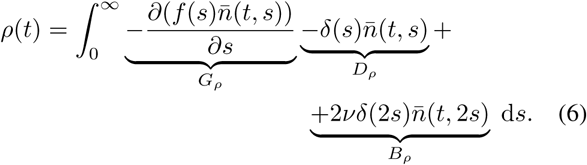

It is important to notice that that *ρ* differs from the average specific growth rate *µ*, normally used to compute the dynamics 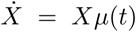, where 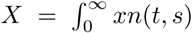 is an extensive property, like size, integrated over the whole population. It should be noted that *ρ* and *µ* coincide only when 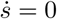 is assumed.

The first term in (6), nullifies. This can be shown by using the product rule on the derivative and integration by parts.

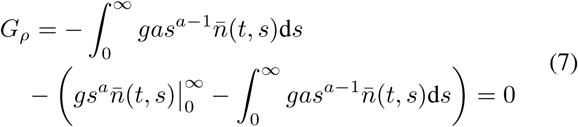

We see that the first and last term in (7) cancel out. We notice that the second term is the integrand function of the *a*^*th*^ raw moment. This goes to zero at +∞, for our previous assumption, and 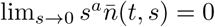, as 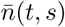 is finite ∀*s*. Using the substitution 2*s* = *z* in the *B*_*ρ*_ term, and noticing that this does not affect the limits of integration, the *B*_*ρ*_ and *D*_*ρ*_ terms can be summed up leaving

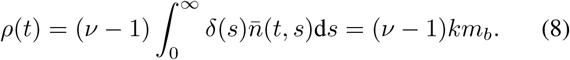

We can already see here that if we choose to model the dynamics of a single lineage, for which *ν* = 1, a value of *ρ*(*t*) = 0 is obtained, as expected.

If we rewrite (1) by replacing 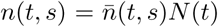 and introduce the expression for the kernel function, we obtain

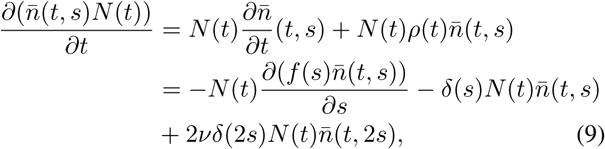

If we cancel *N* (*t*) from both sides, the normalized PBE can be obtained as

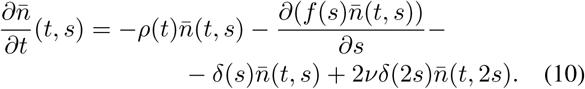

Taking the *i*^*th*^ moment *m*_*i*_, for *i* ≥ 1, of the normalized PBE (10) results in

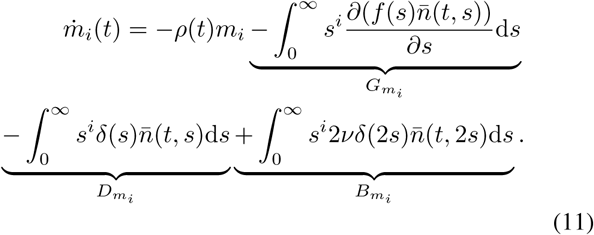

Applying the product rule followed by integration by parts, and exploiting our assumptions, as we did in (7), the term 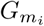 resolves in

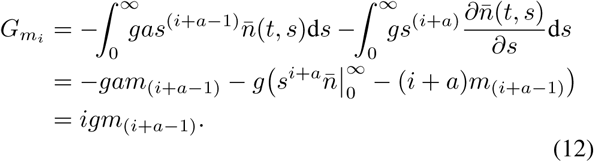

After the substitution 2*s* = *z* in 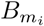, the terms 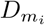 and 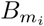 can be summed up, leaving us with

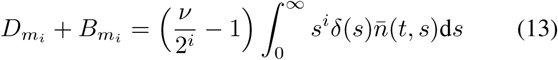

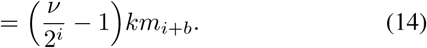

From equations (12) and (14) we notice that, for *a >* 1 or *b >* 0, the moment dynamics are not closed. This dependency is overcome through the moment closure method [13], for which all central moments higher than *cm*_*i*_ are approximated to zero. Introducing *p* = max((*a* − 1), *b*) we write:

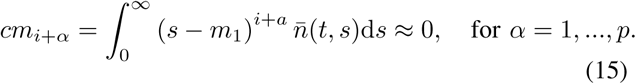

This approximation offers us algebraic expressions to compute higher moments, and thus the term in (12) and (14). As an example, the approximated expressions for *m*_*i*+*α*_ when *i* = 2, 3 and *p* = 1, are provided here below:

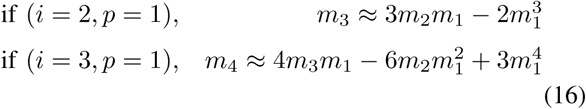

Putting all together, (11) becomes:

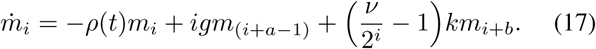

#### b) Stochastic Simulation Algorithm (SSA)

The Stochastic Simulation Algorithm is a numerical approach for generating sample paths of a continuous-time Markov process whose probability distribution evolves through transition rates between its states. For each cell *i* in the population, the algorithm first computes *P*_*i*_(*t*), the probability that cell *i* divides at time *t*, by integrating the division rate *δ* over time (see explanation in Supplementary Information)[12]. Then, these *P*_*i*_(*t*) are compared to uniformly generated random numbers *r*_*i*_, and the next division is implemented only for cell *j*, with *P*_*j*_ *> r*_*j*_. The algorithm is detailed as follows:

##### Algorithm 1: Stochastic Simulation Algorithm

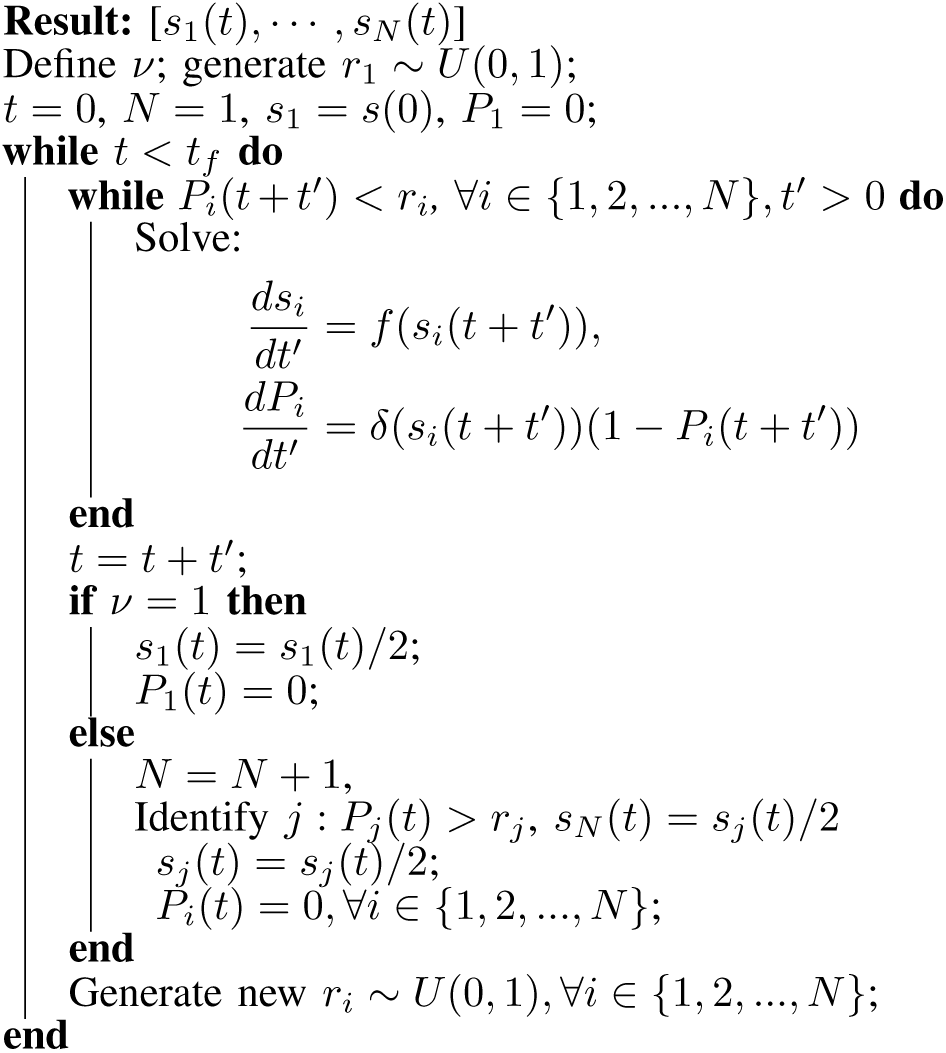

Giving us the sizes [*s*_1_ *… s*_*N*_] of the cells at the time *t* ∈ (0, *t*_*f*_) for each replica. The single-cell growth rate *f* (*s*), the division rate *δ*(*s*) and *ν*, the number of cells tracked after division, are used consistently with their definition in the PBE (see (1)). Let *L* be the number of replicas, *N*_*l*_(*t*) the number of cells in replica *l* at time *t*, 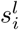 the size of cell *i* in replica *l*. The mean population number *N* (*t*), the statistical mean ⟨*s*⟩ = *m*_1_ and variance 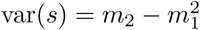 of the cell size distribution are computed as follows:

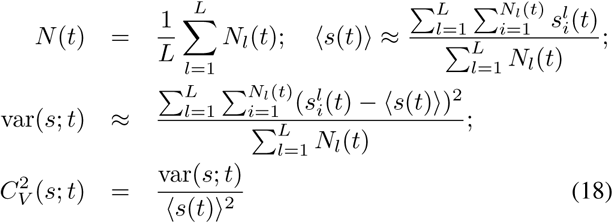

In this work we make use of the squared coefficient of variation 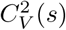, a dimensionless quantity that allows us to estimate the degree of heterogeneity in the distribution. For ease of notation, in the rest of the paper the 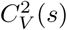 of the distribution of sizes *s* is replaced by 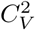.

## III. RESULTS AND DISCUSSION

### A. Biological scenarios

For our analysis we define four biologically relevant scenarios, that differ for the cell growth regime and division mechanisms assumed. The single-cell growth rate *f* is thus defined as either *f* (*s*) = *g* (linear growth), or *f* (*s*) = *gs* (exponential growth). Similarly, the division rate *d* is considered either constant, *δ*(*s*) = *k*, or linear, *δ*(*s*) = *ks*, with respect to size.

For each scenario, the parameter values for *g* and *k* shown in table I were chosen so that the mean doubling time *τ* satisfies *τ* = 1, and 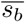, defined the mean size at birth, satisfies 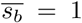. This 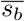, introduced in previous studies[12] (see Supplementary Information), corresponds to the size that is perfectly doubled when division occurs. Importantly, specific values for both *τ* and 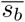 can be determined experimentally, so that experimental data, once normalized to these values, can be compared to model predictions.

**TABLE I.** Description of the scenarios defined by their growth regime *f* and division mechanism *d*.

| Sc. | $p$ | $a$ | $f(s)$ | $\delta(s)$ | $\rho$ |
| --- | --- | --- | --- | --- | --- |
| I | 0 | 0 | $g = 1$ | $k = 1$ | $\nu - 1$ |
| II | 1 | 0 | $gs = \ln(2)s$ | $k = 2 \ln(2)$ | $\nu - 1$ |
| III | 0 | 1 | $g = 1$ | $ks = 0.562s$ | $(\nu - 1)0.562m_1$ |
| IV | 1 | 1 | $gs = \ln(2)s$ | $ks = \ln(2)s$ | $(\nu - 1) \ln(2)m_1$ |

### B. Computational results

Here, we compare the results of the two approaches presented in II, with respect to the dynamics of the mean size ⟨*s*⟩ and of the coefficient of variation 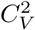. The initial size distribution was defined as a Dirac-Delta with either 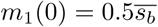 or 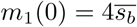. These two initial values can be considered as the extremes of a range of biologically realistic cell sizes. All the higher moments have initial conditions *m*_*i*_(0) = (*m*_1_(0))^*i*^. The initial number of cells in the population is set to *N* (0) = 1.

#### a) Population Balance Equation

The analytical expressions derived from the general PBE, presented in Section II, are used to compute the dynamics for *m*_1_ and *m*_2_, and either *ν* = 1 or *ν* = 2, and reported in table II. For *b* = 1, the moment closure approximation is used and *cm*_3_ ≈ 0.

**TABLE II.** Dynamics of the first and second raw moments using the moment closure approximation at the third moment.

| | $\nu = 1$ | $\nu = 2$ |
| --- | --- | --- |
| Sc | Moment dynamics | Moment dynamics |
| I | $\dot{m}_1 = g - \frac{1}{2}km_1$<br>$\dot{m}_2 = 2gm_1 - \frac{3}{4}km_2$ | $\dot{m}_1 = g - km_1$<br>$\dot{m}_2 = 2gm_1 - \frac{3}{2}km_2$ |
| II | $\dot{m}_1 = gm_1 - \frac{1}{2}km_1$<br>$\dot{m}_2 = (2g - \frac{3}{4}k)m_2$ | $\dot{m}_1 = (g - k)m_1$<br>$\dot{m}_2 = (2g - \frac{3}{2}k)m_2$ |
| III | $\dot{m}_1 = g - \frac{1}{2}km_2$<br>$\dot{m}_2 = 2gm_1 + \frac{3}{2}km_1^3 - \frac{9}{4}km_1m_2$ | $\dot{m}_1 = g - km_1^2$<br>$\dot{m}_2 = 2gm_1 + km_1^3 - \frac{5}{2}km_1m_2$ |
| IV | $\dot{m}_1 = gm_1 - \frac{1}{2}km_2$<br>$\dot{m}_2 = 2gm_2 + \frac{3}{2}km_1^3 - \frac{9}{4}km_1m_2$ | $\dot{m}_1 = gm_1 - km_1^2$<br>$\dot{m}_2 = 2gm_2 + km_1^3 - \frac{5}{2}km_1m_2$ |
| | Equilibrium points $[\tilde{m}_1, \tilde{m}_2]$ | Equilibrium points $[\tilde{m}_1, \tilde{m}_2]$ |
| I | $[\frac{2g}{k}, \frac{16g^2}{3k^2}]$, stable $\forall g, k > 0$ | $[\frac{g}{k}, \frac{4g^2}{3k^2}]$, stable $\forall g, k > 0$ |
| II | $[0, 0]$, stable if $g < \frac{3k}{8}$ | $[0, 0]$, stable if $g < \frac{3k}{4}$ |
| III | $[\sqrt{\frac{5g}{3k}}, \frac{2g}{k}]$, stable $\forall g, k > 0$<br>$[0, \frac{2g}{k}]$, unstable $\forall g, k > 0$ | $[\sqrt{\frac{g}{k}}, \frac{6g}{5k}]$, stable $\forall g, k > 0$ |
| IV | $[0, 0]$, unstable $\forall g, k > 0$ | $[\frac{g}{k}, \frac{2g^2}{k^2}]$, stable $\forall g, k > 0$<br>$[0, 0]$, unstable $\forall g, k > 0$ |

The PBE is here solved also numerically, and the moment dynamics are calculated from the dynamics of the full distribution. This offers us a comparison to the results obtained through the analytical derivations. In the literature, different numerical schemes have been proposed to solve PBEs. We exploited the cell average technique, a spatial discretization method that, following the implementation in [14], preserves the first two moments *m*_0_ and *m*_1_ of the distribution.

#### b) Stochastic Simulation Algorithm (SSA)

To study the behavior of single lineages (*ν* = 1), the dynamics of 20K replicas are simulated. To study the population snapshot perspective (*ν* = 2), 1K replicas are run, all starting from one cell with the same initial size (*s*(0)). Growth and division rates are defined according to the specific scenarios and the values of *g* and *k*. At the predefined time points the mean size ⟨*s*⟩ and the coefficient of variation 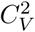 are computed considering the sizes of all cells across all replicas using expressions (18). To integrate the differential equations in Algorithm 1, we use Euler integration.

Figures 1-4 show the resulting dynamics of mean size ⟨*s*⟩, in units of 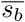, and the variability in terms of 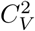. In figure 5 we report the comparison of the dynamics of the total number of cells in the simulation.

**Fig. 1.**
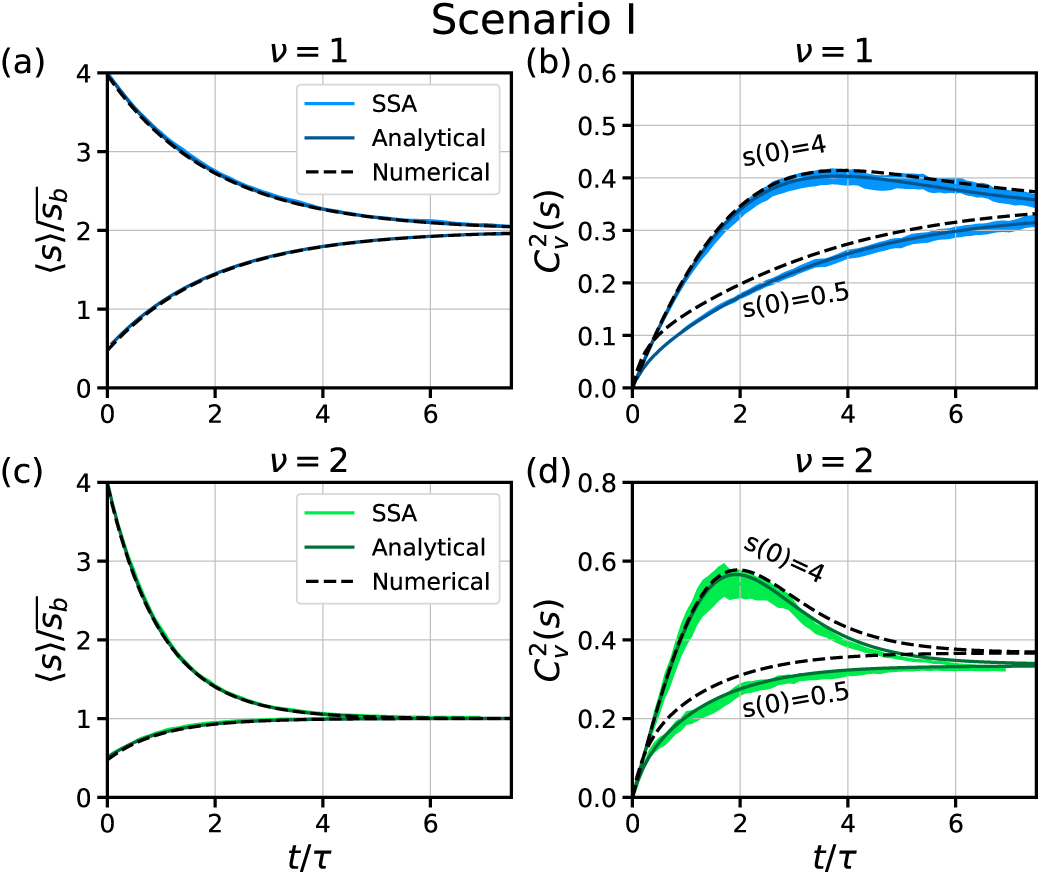
Size dynamics for cells growing and dividing in the scenario I (*f* = *g* = 1, *b* = *k* = 1). Analytical, Numerical and stochastic simulation (SSA) are compared. (a) Mean size ⟨*s*⟩ dynamics (b) Size variability dynamics 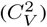. Single lineage is considered (*ν* = 1) for both (a) and (b). (c) Mean size dynamics (d) Size variability. Population snapshot is considered (*ν* = 2) or both (c) and (d). Width of the line of SSA represents the 95% confidence interval for 20K cell for *ν* = 1 and 1*K* population replicas for *ν* = 2. Two different initial conditions (*m*_1_(0) = 4 and *m*_1_(0) = 0.5) was considered in all cases.

**Fig. 2.**
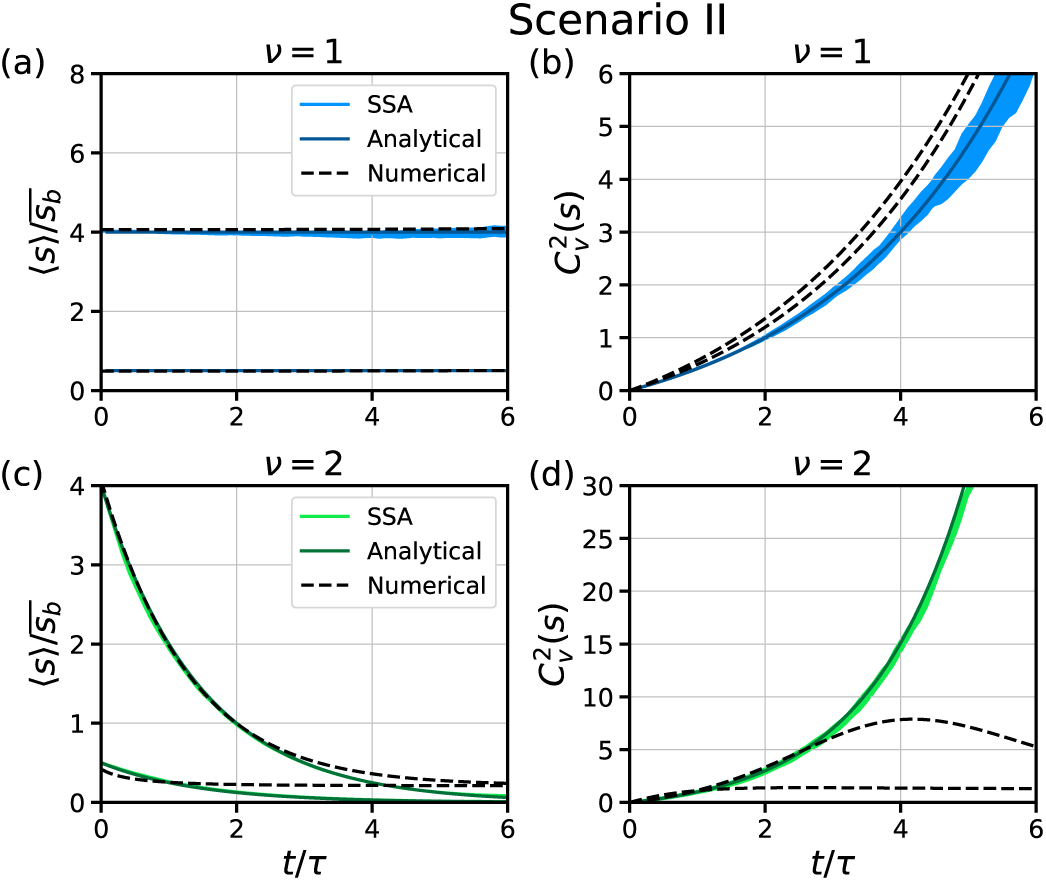
Size dynamics for cells growing and dividing in the scenario II (*f* = *gs* = ln(2)*s, b* = *k* = 2 ln(2)). Analytical, Numerical and stochastic simulation (SSA) are compared. (a) Mean size ⟨*s*⟩ dynamics (b) Size variability dynamics 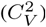. Single lineage is considered (*ν* = 1) for both (a) and (b). (c) Mean size dynamics (d) Size variability. Population snapshot is considered (*ν* = 2) or both (c) and (d). Width of the line of SSA represents the 95% confidence interval for 20K cell for *ν* = 1 and 1*K* population replicas for *ν* = 2. Two different initial conditions (*m*_1_(0) = 4 and *m*_1_(0) = 0.5) was considered in all cases.

**Fig. 3.**
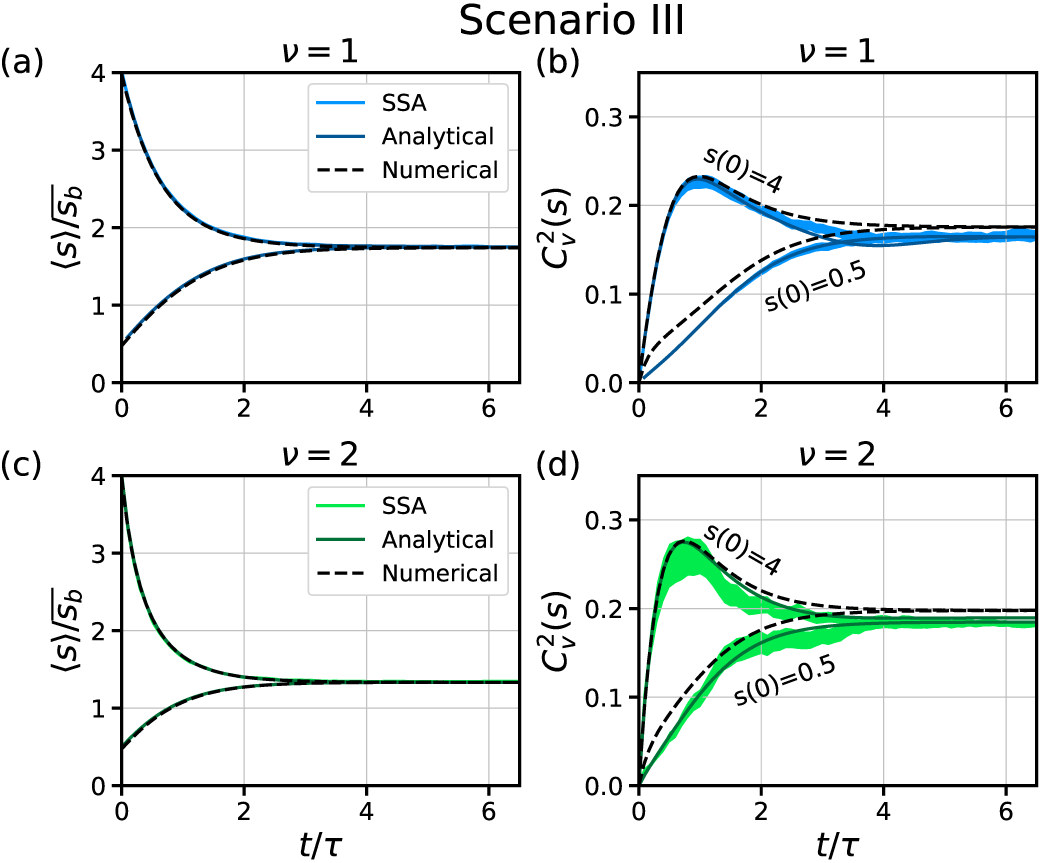
Size dynamics for cells growing and dividing in the scenario III (*f* = *g* = 1, *b* = *ks* = 0.562*s*). Analytical, Numerical and stochastic simulation (SSA) are compared. (a) Mean size ⟨*s*⟩ dynamics (b) Size variability dynamics 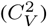. Single lineage is considered (*ν* = 1) for both (a) and (b). (c) Mean size dynamics (d) Size variability. Population snapshot is considered (*ν* = 2) or both (c) and (d). Width of the line of SSA represents the 95% confidence interval for 20K cell for *ν* = 1 and 1*K* population replicas for *ν* = 2. Two different initial conditions (*m*_1_(0) = 4 and *m*_1_(0) = 0.5) was considered in all cases.

**Fig. 4.**
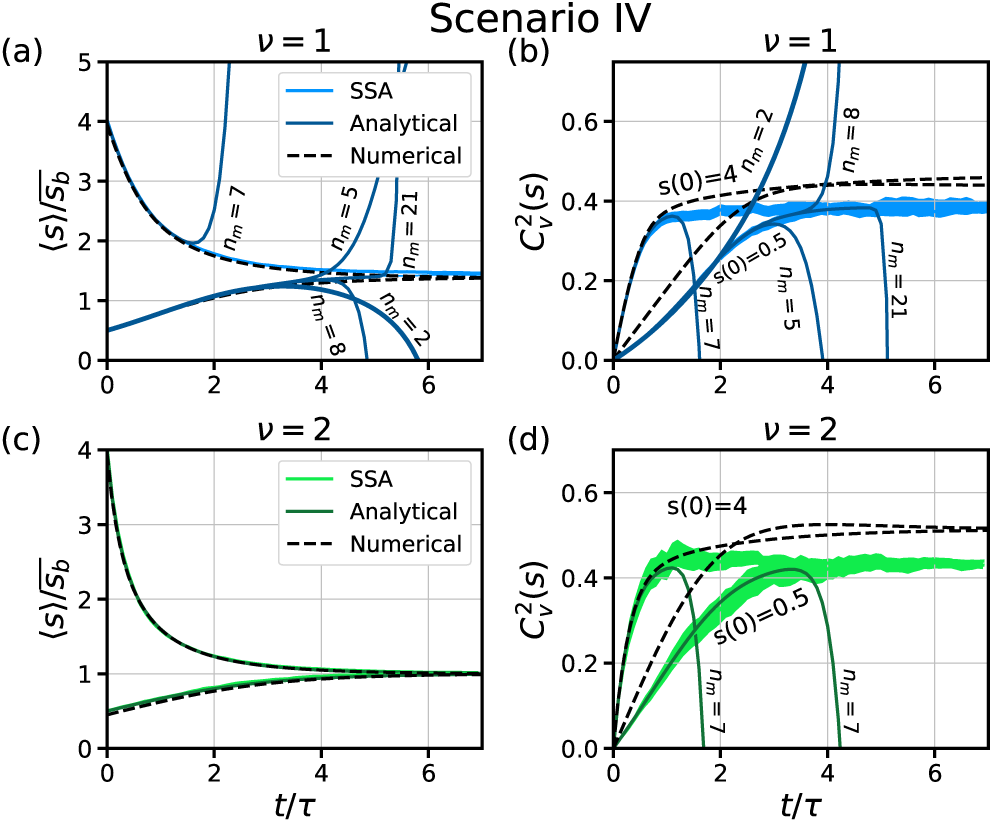
Size dynamics for cells growing and dividing in the scenario IV (*f* = *gs* = ln(2)*s, b* = *ks* = ln(2)*s*). Analytical, Numerical and stochastic simulation (SSA) are compared. ((a) Mean size ⟨*s*⟩ dynamics (b) Size variability dynamics 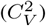. Single lineage is considered (*ν* = 1) for both (a) and (b). Different levels (*nm*) of ODE truncation using moment closure are shown in (a) and (b). (c) Mean size dynamics (d) Size variability. Population snapshot is considered (*ν* = 2) or both (c) and (d). Width of the line of SSA represents the 95% confidence interval for 20K cell for *ν* = 1 and 1*K* population replicas for *ν* = 2. Two different initial conditions (*m*_1_(0) = 4 and *m*_1_(0) = 0.5) was considered in all cases.

**Fig. 5.**
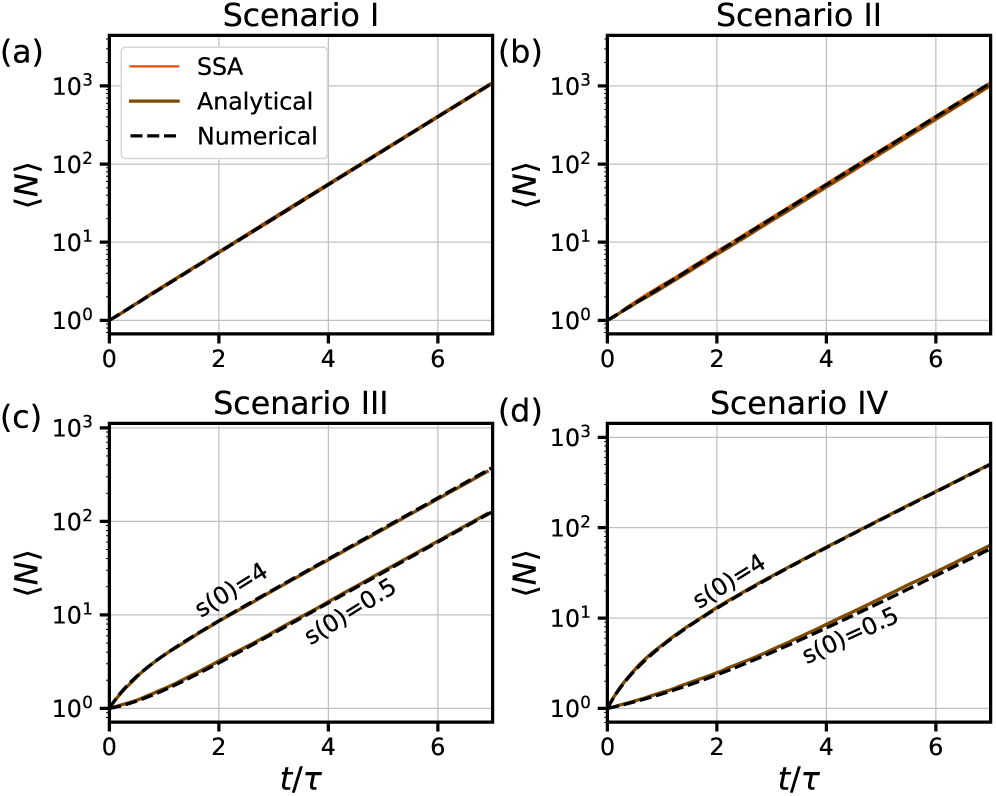
Total number of particles in the population. Scenarios 1-4 are labeled respectively. Width of the line of SSA represents the 95% confidence interval for 1*K* population replicas.

### C. Discussion

Cell size heterogeneity can be quantified across single lineages or taking snapshots of the whole population. This crucial difference is represented, in the equations shown so far, by the value of *ν*. Depending on the organism and the availability of extracellular nutrients, cells can adopt different growth regimes and division strategies. Considering these, we defined the four scenarios presented in table I.

Figures 1 - 4 show that the four scenarios result in clearly distinct dynamics for ⟨*s*⟩ and 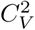. Taking into consideration all three trajectories, we see that in scenarios I, III and IV, a stable steady state is reached. In scenario II, instead, no stable steady state is reached, confirming the analytical results in table II and findings reported in other studies [15]. It is fair to consider that this scenario is not realistic, as cells with excessively different sizes are generated.

Transient dynamics reveal that, for all scenarios and *ν* values, the steady state values 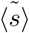 and 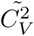 do not depend on the initial conditions.

The comparison between single lineages (*ν* = 1) and population snapshots (*ν* = 2), shows us that the computed steady-state values 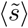 are consistently higher for single lineages and lower for population snapshots. As reported in similar studies [6], this is expected, because, independently of the division mechanism, cells that proliferate faster are on average smaller and over-represented in the whole population. Moving one step further this observation, we argue that our computational framework and our results can be used to connect the moment dynamics in the two perspectives we examined. More clearly, if, for instance, one of the scenarios in table I can be considered an accurate representation of an experiment on cell lineages, then, exploiting our framework, the data collected from cell lineages could be used to infer the dynamics of population snapshot. Vice versa, data from population snapshots could be used to infer the dynamics in single lineages. For scenarios I, II and III, the analytical formulas in table II would make the inference straightforward. For scenario IV, where the analytical dynamics are inaccurate, one would need to first use the numerical approach or the SSA to infer *g* and *k* parameters and then, after changing the value of *ν*, use these to simulate the unknown dynamics.

Despite the clear differences in 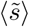, it is important to notice that the steady-state values 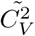, when finite, do not show appreciable differences when different *ν*s are considered. Overall, we can conclude that experimentally, different average sizes, but not significantly different variabilities should be expected for single lineages versus population snapshots.

The use of different modeling approaches and solution schemes offers us an indication on the accuracy of the results: if two trajectories out of three superimpose, their accuracy is reinforced. In scenarios I and II, no moment closure approximation is introduced, thus the analytical solutions are considered as the correct dynamics. As the stochastic simulations lie in close proximity to the analytical curves, we consider that the SSA is accurate. A similar closeness is found again in scenario III, where, however, the analytical moment dynamics are not closed. We can thus conclude that the moment closure approximation [13] is here accurate. In scenario IV, instead, the approximation is unreliable. From Figures 4 (a) and 4 (b) we see that, even if we increase significantly the order of the moment (*n*_*m*_) where the moment-closure approximation is applied, the accuracy of the analytic trajectory is preserved only for a limited time. In this case, SSA predictions are corroborated by the numerical solution. Implementing the cell average technique as in [14], the accuracy of numerical results is only guaranteed for 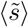 dynamics. This approach looses precision, however, when the 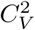 rapidly diverges, like in scenario II, as the discretization grid can no more cover the higher cell sizes in the population. Despite these limitations, the numerical solution of the PBE offers a useful comparison at a much lower computational cost than the stochastic approach.

Figure 5 shows that, for constant division rates, the exponential increase in *N*, the total number of cells in the population, is unaffected by the initial condition. For size-dependent division rates, instead, if 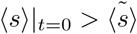 the population starts expanding rapidly, while, if 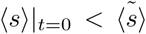, the expansion displays a small lag. In both cases an exponential regime is reached after a few doubling times.

## IV. CONCLUSIONS

Cell size heterogeneity in growing populations is tightly controlled and generally low, a phenomenon defined as size homeostasis. In this work, we used multiple mathematical modeling approaches to provide a further understanding on how studies on size homeostasis are influenced by changes in different model components. On one side, we considered different combinations of growth regimes and division strategies, which lead us to define four different biologically relevant scenarios. On the other, we studies the variability of cell sizes in a population observed from two perspectives: single lineages and population snapshots.

We used a deterministic population-based framework, the population balance equation (PBE), which models growth and division process trough continuous rates. This approach allowed us to derive analytically a system of ordinary differential equations (ODE) that describes the dynamics of the raw moments of the cell size distribution. For some of the considered scenarios, the moment closure approximation was used to truncate the system and avoid dependencies on higher-order moments. The solution of this ODE system was compared with a numerical solution of the PBE and with stochastic simulations. Our comparison showed that the analytical derivations offer a good approximation in three out of four scenarios.

Secondly, the mean cell-size for single lineages is consistently higher than for population snapshots, while their variability 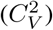 is not significantly different. In some of the scenarios, we show that the ODEs allow us to establish mathematical relationships among the two considered perspectives. With this, we propose that the steady state values for population snapshots can be inferred from results or data on single lineages, and vice versa.

## V. ACKNOWLEDGMENTS

We thank Juan Manuel Pedraza at Universidad de los Andes for the discussion.

## Notes

### Competing Interest Statement

The authors have declared no competing interest.

