## Supplemental information for "Cell size statistics in cell lineages and population snapshots with different growth regimes and division strategies"

#### S1 The division strategies in bacteria

Following the literature, the strategy by which a bacterial cell divides can be classified into three main strategies or paradigms. These are shown in Figure S1, where an exponential growth regime is assumed. The first strategy is the *timer* (Fig. S1(a)), in which cells wait a fixed time, on average, and then divide. The second is the *adder* (Fig. S1(b)), in which cells attempt to add a fixed size, on average, before dividing. The third is the *sizer* (Fig. S1(c)), in which cells grow until a certain volume is reached.

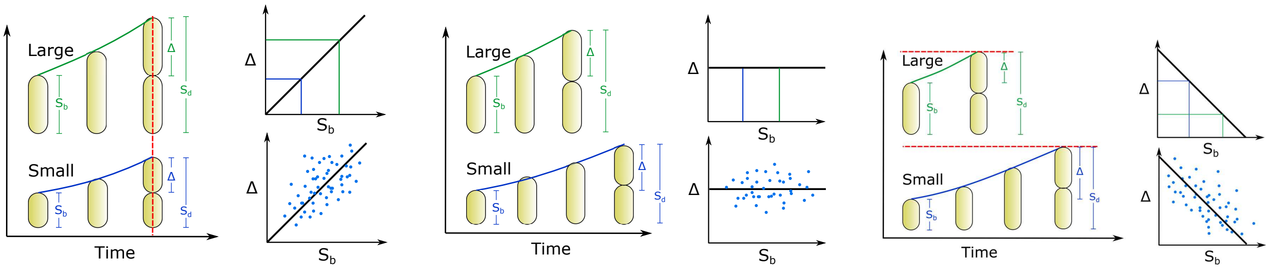


(a) Timer strategy: Cell waits a fixed time before splitting.

(b) Adder strategy: Cell adds a fixed size before splitting.

(c) Sizer strategy: Cell waits to reach a given size before splitting.

Figure S1: Main division strategies for exponentially growing bacteria. The size at division is shown for to starting sizes: one large and other small showing the differences in added size. The plot of $\Delta$ vs $s_b$ is also shown for all the strategies.

These strategies can be distinguished by observing the trend line that describes the relationship between the added size, $\Delta$, versus the size at birth, $s_b$[1]. The *timer* strategy shows a positive correlation between them: the bigger the size at birth, the bigger the cell at division, as Figure 1(a) shows. When cells follow the *adder* strategy, instead, they gain a constant added size and their size at division does not depend on the size at birth. The *sizer* paradigm, in the end, produces a slope of -1, given the relationship between the size at birth and the size that needs to be added to reach the desired fixed size at division. It is often the case, however, that intermediate values for the slopes are observed. We thus define the sizer-like strategy, for a negative slope between $-1$ and 0, and the timer-like strategy, for a positive slope between 0 and 1.

#### S2 Estimation of division properties for the studied scenarios

In the main text we defined that the division rate $\delta(s(t))$ is dependent on time implicitly through the size $s(t)$, since cells grow along the time. If we consider $t$ as the time spent by the cell in the cell cycle since its birth, we can represent division as a single step stochastic process, with occurrence rate $\delta(t)$, between the state "0" (not divided yet) to the state "1" (already divided). The probability, or cumulative probability function, $P_i$ of being in state $i$ changes in time according to the following master equation:

$$\begin{aligned} \frac{dP_0}{dt} &= -\delta(t)P_0(t) \\ \frac{dP_1}{dt} &= \delta(t)P_0(t), \end{aligned} \quad (1)$$

which, assuming boundary conditions $P_0(t=0)=1$, $P_1(t=0)=0$, has the solution:

$$P_0(t) = \exp\left(-\int_0^t \delta(t')dt'\right)$$
$$P_1(t) = 1-\exp\left(-\int_0^t \delta(t')dt'\right). \tag{2}$$

Recalling that $P_1(t)$ corresponds precisely to the cumulative probability of being divided at time $t$, we can simplify its notation to $P(t)$. Then, reintroducing the explicit dependency $\delta(t') = \delta(s(t'))$ we get:

$$P(t) = 1-\exp\left(-\int_0^t \delta(s(t'))dt'\right). \tag{3}$$

In the main text we presented four different scenarios, which differ for the growth regime and the division strategy assumed. If we define $s_0$ as the cell size at an arbitrary time $t_0$, the division rate $\delta$ can be computed through the formulas in Table S1:

| Scenario | $\delta(s(t))$ |
|---|---|
| I | $k$ |
| II | $k$ |
| III | $ks(t) = k(s_0 + g(t-t_0))$ |
| IV | $ks(t) = k(s_0 \exp[g(t-t_0)])$ |

Table S1: Division rates as function of time.

the inverse problem, this is, to find the division rate $\delta(s(t))$ knowing the probability of being divided $P(t)$, can be solved from (3):

$$\delta(s(t)) = -\frac{d}{dt}\left[\ln\left(1-P(t)\right)\right] = \frac{1}{(1-P(t))}\frac{dP(t)}{dt}. \tag{4}$$

Since (4) is equivalent to (3), we choose (4) to be solved in the Stochastic Simulation Algorithm.

Deriving $P(t)$ with respect to $t$, we get the probability density function $\rho(t)$ of time at division

$$\rho(t) = \frac{dP(t)}{dt}, \tag{5}$$

and then the distribution of size at division $\rho(s_d)$ by using a transformation of variables

$$\rho(s_d) = \rho(t(s_d))\frac{dt}{ds_d}, \tag{6}$$

where $\frac{dt}{ds_d}$ can change depending on the growth regime. We consider the general case where $\rho(s_d)$ depends on the cell size at birth $s_b$, which can be a random variable as well. Thus, we redefine $\rho(s_d)$ as a conditional probability $\rho(s_d|s_b)$.

Thus, the average size at division can be estimated:

$$\langle s_d\rangle \equiv \int_{s_b}^{\infty} s_d \rho(s_d|s_b) ds_d, \tag{7}$$

where $\rho(s_d|s_b)$ is the conditional distribution of size at division $s_d$ given the size at birth $s_b$, obtained using (6).

### S3 The definition of $\overline{s_b}$

In the main article, we introduced $\overline{s_b}$, as the unit of cell size. This corresponds to the mean size at birth, and, because we assumed equally-sized daughter cells, it is also the size that is perfectly doubled when division occurs.

In order to explain why this size unit is so important, we make the example of a generic bacterium following the adder strategy, which implies that its size at division $s_d$ can be defined as $\langle s_d\rangle = s_b + \Delta$, with $\Delta$ some constant. Independently of its starting size $s_0$, if we wait that enough cell cycles are completed, we will observe that the size of its progeny approaches $\overline{s_b}$. This phenomenon, termed as size convergence, has been widely observed in experiments [2]. Defining $s_i$ the size of the newborn cell after the $i^{th}$ division, we get:

$$\begin{aligned} s_1 &= \frac{s_0+\Delta}{2} \\ s_2 &= \frac{s_1+\Delta}{2} = \frac{s_0+3\Delta}{4} \\ s_n &= \frac{s_0+(2^n-1)\Delta}{2^n} \\ \overline{s_b} &= \lim_{n\to=\infty} s_n = \Delta. \end{aligned} \tag{8}$$

Figure S2 portrays this iterative process, and shows how the sizes at birth relate, on one side, to the adder division law, represented by the green dashed line, and, on the other side, to the assumption of symmetric division ($s_b + \frac{s_d}{2}$), represented by the blue solid line. The pink line describes the evolution of sizes across multiple generations. For $\lim_{n\to\infty}$, we can see that its trajectory converges to the point where $s_d = 2s_b = s_b + \Delta$. For the adder division strategy, this is the size that satisfies $\overline{s_b} = \Delta$.
So far in the explanation we assumed an adder strategy. If the timer or sizer paradigms are instead considered, $\overline{s_b}$ can be estimated numerically by solving (7):

$$\overline{s_b} \equiv s_b : \langle s_d\rangle(s_b) = 2s_b, \tag{9}$$

that is, $\overline{s_b}$ is the size at birth $s_b$ such as its mean size at division $\langle s_d\rangle$ is twice itself $2s_b$.

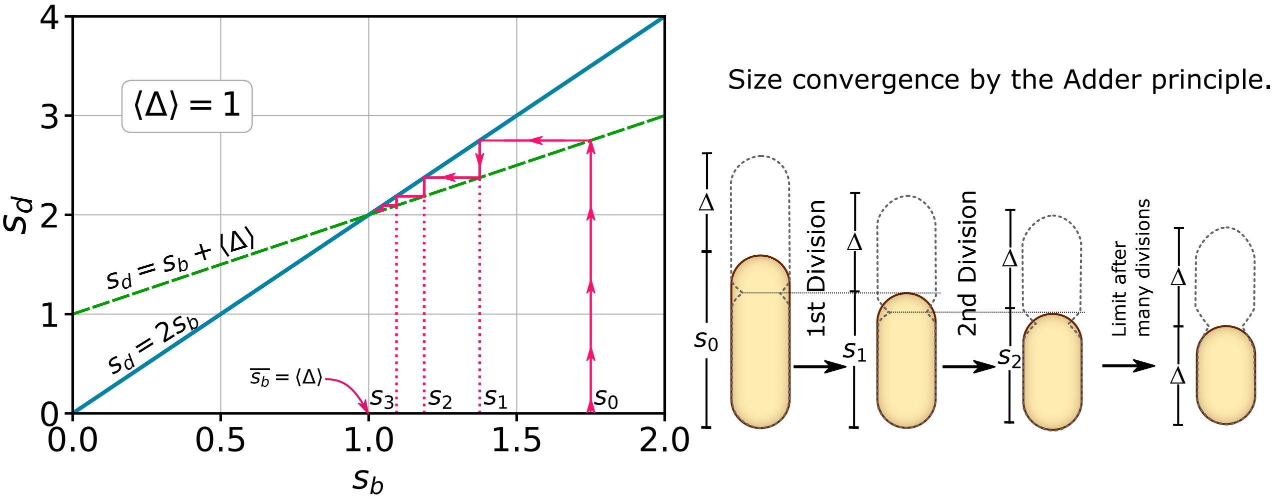


Figure S2: Convergence of the cell size to $\overline{s_b}$ explained for the adder strategy. After enough divisions are completed, the size at birth converges to $\overline{s_b}$, satisfying $\overline{s_b} + \Delta = 2\overline{s_b}$, and thus $\overline{s_b} = \Delta$.

If we connect the concepts introduced so far, we see that a division strategy is classified as timer, adder or sizer depending on $\rho(s_d|s_b)$, and how it affects $\langle s_d\rangle(s_b)$ in (7), and depending on the slope in the relationshp between $\Delta$ vs $s_b$, depicted in figure S1.

### S4 Statistical properties of the division scenarios

In this work, as we mention in the main text, in order to obtain results that are comparable to experiments, we fixed $k$ and $g$ so that $\bar{s_b} = 1$, and all sizes are normalized to it. The average size at birth $\bar{s_b}$, in fact, represents a parameter that can be measured experimentally. The values of $k$ and $g$ satisfying $\bar{s_b} = 1$ can be obtained by solving (9) either analytically o numerically.

To define our parameters, we impose another constraint on the doubling time $\tau$, the time needed for a cell whose size at birth is $s_b = 1$ to double its size, so that $\tau = 1$. For a linear growth regime, this results into $g = 1$, while for exponential growth, it produces $g = \ln(2)$.

We list here the statistical properties that can be obtained analytically using the framework explained in this work: the growth regime, the division rate, the constants $k$ (division constant) and $g$ (growth rate), the geometric transformation differential $\frac{dt}{ds_d}$ used in (6), the distribution of the time to division $\rho(t)$ since cell birth, the distribution $\rho(s_d|s_b)$ of size at division $s_d$ given the size at birth $s_b$. All of these properties are summarized in tables S2 and S2.

| Properties | Scenario I | Scenario II |
|---|---|---|
| Growth Regime | $s = s_b + gt$ | $s = s_b \exp(gt)$ |
| Division Rate | $k$ | $k$ |
| $k$ | 1 | 2ln(2) |
| $g$ | 1 | ln(2) |
| $\frac{dt}{ds_d}$ | $\frac{1}{g}$ | $\frac{1}{gs_d}$ |
| $\rho(t)$ | $k\exp(-kt)$ | $k\exp(-kt)$ |
| $\rho(s_d\|s_b)$ | $\frac{k}{g}\exp\left(-k\frac{s_d-s_b}{g}\right)\theta(s_d-s_b)$ | $\frac{k}{g}\left(\frac{s_b}{s_d}\right)^{\frac{k}{g}}\frac{1}{s_d}\theta(s_d-s_b)$ |
| Division Law | $\langle s_d\rangle = s_b + \frac{g}{k}$ | $\langle s_d\rangle = s_b + s_b\left(\frac{1}{1-\frac{g}{k}}\right)$ |
| Division Strategy | *Adder* | *Timer* |

Table S2: Statistic properties for the scenarios I and II.

| Properties | Scenario III | Scenario IV |
|---|---|---|
| Growth Regime | $s = s_b + gt$ | $s = s_b \exp(gt)$ |
| Division Rate | $ks$ | $ks$ |
| $k$ | 0.562 | ln(2) |
| $g$ | 1 | ln(2) |
| $\frac{dt}{ds_d}$ | $\frac{1}{g}$ | $\frac{1}{gs_d}$ |
| $\rho(t)$ | $k(s_b+gt)\exp\left[-k\left(s_bt+g\frac{t^2}{2}\right)\right]$ | $ks_b\exp\left[-\frac{k}{g}s_b\left(\exp(gt)-1\right)+gt\right]$ |
| $\rho(s_d\|s_b)$ | $\frac{k}{g}s_d\exp\left[-\frac{k}{g}\left(\frac{s_d^2-s_b^2}{2}\right)\right]\theta(s_d-s_b)$ | $\frac{k}{g}\exp\left[-\frac{k}{g}(s_d-s_b)\right]\theta(s_d-s_b)$ |
| Division Law | $\langle s_d\rangle = s_b + \sqrt{\left(\frac{g}{k}\right)}\exp\left(\frac{k}{g}\frac{s_b^2}{2}\right)\sqrt{\frac{\pi}{2}}\mathrm{erf}\left(\frac{s_b}{\sqrt{2(g/k)}}\right)$ | $\langle s_d\rangle = s_b + \frac{g}{k}$ |
| Division Strategy | *Sizer-like* | *Adder* |

Table S3: Statistic properties for the scenarios III and IV. $\theta(x)$ is the Heaviside step function, erf(x) $= \frac{2}{\sqrt{\pi}}\int_0^x e^{-t^2}dt$.
